# The FXR2P low complexity domain drives assembly of multiple fibril types with differing ribosome association in neurons

**DOI:** 10.1101/505255

**Authors:** Emily E. Stackpole, Michael R. Akins, Maria Ivshina, Anastasia C. Murthy, Nicolas L. Fawzi, Justin R. Fallon

## Abstract

RNA binding proteins (RBPs) typically function in higher order assemblages to regulate RNA localization and translation. The Fragile X homolog FXR2P is an RBP essential for formation of Fragile X granules, which associate with axonal mRNA and ribosomes in the intact brain. Here we performed an unbiased EGFP insertional mutagenesis screen to probe for FXR2P domains important for assembly into higher order structural states in neurons. Fifteen of the 18 unique in-frame FXR2P^EGFP^ fusions tested formed cytosolic granules. However, EGFP insertion within a 23 amino acid region of the low complexity (LC) domain induced formation of distinct FXR2P^EGFP^ fibrils (A and B) that were found in isolation or assembled into highly ordered bundles. Type A and B complexes exhibited different developmental timelines, ultrastructure and ribosome association with ribosomes absent from bundled Type B fibrils. The formation of both fibril types was dependent on an intact RNA binding domain. We conclude that formation of these higher order FXR2P assemblages with alternative structural and compositional states in neurons requires collaboration between the LC and RNA binding domains.

**Summary Statement:** The Fragile X protein FXR2P forms multiple types of fibrillar assemblages with differential ribosome associations in neurons through cooperation between its RNA binding and LC domains.

## INTRODUCTION

Neurons are highly elaborate cells that mount specific and dynamic responses to a range of stimuli within their vast subdomains. One important mechanism for such spatiotemporal control is local protein synthesis where mRNAs are targeted to specific domains and can be rapidly translated in response to nearby cues. This process has long been appreciated in dendrites (Holt and Schuman 2013; Sutton and Schuman 2006) and recent evidence indicates that analogous translational machinery is also present in axons (Zheng et al., 2001; Christie et al., 2009; Taylor et al., 2009; Korsak et al., 2016; Shigeoka et al., 2016; Akins et al., 2017; Batista et al., 2017). Local protein synthesis is in part controlled by RNA binding proteins (RBPs) that can influence transcript fate through regulation of RNA biogenesis, localization, translation and degradation (Kapeli & Yeo, 2012; Darnell, 2013; Calabretta and Richard, 2015). Further, RBP alteration or loss is the basis for a wide range of neurological diseases including Fragile X syndrome, frontotemporal dementia, spinal muscular atrophy, myotonic dystrophy and amyotrophic lateral sclerosis (Liu-Yesucevitz et al., 2011; Wang et al., 2016).

RBPs are integral components of RNA granules, a class of cytosolic ‘assemblages’ that are critical regulators of mRNA transport, targeting and local translation (Kiebler and Bassell, 2006; Toretsky and Wright, 2014; Buchan, 2014; Calabretta and Richard, 2015; Nielsen et al., 2016). Formation of RNA granules is dependent upon the low complexity (LC) domains that are present in a majority of RBPs (Kato et al., 2012; Han et al., 2012; Weber and Brangwynne 2012; King et al., 2012; Guo and Shorter 2015). LC domains are intrinsically disordered but can promote self-association to collect proteins into highly concentrated states, possibly via changes in LC conformation (Tompa, 2012; Boke et al., 2016; Boeynaems et al, 2018). Therefore, some RBPs harboring LC domains are capable of existing in multiple structural conformations - with an ordered configuration presumably underlying a unique functional state. For example, Xvelo can assemble into an amyloid-like state in the Balbiani body in *Xenopus* (Boke et al., 2016).

RBP-RNA associations are a driving force behind LC domain fibrillization and complex macromolecular organization (Weber & Brangwynne, 2012; Schwartz et al., 2013; Burke et al., 2015; Molliex et al., 2015; Elbaum-Garfinkle et al., 2015). The importance of the strict regulation of the structural states of RBPs is highlighted by the large number of LC domain mutations that cause neurodegenerative diseases (Kim et al., 2013; Maziuk et al., 2017; Harrison & Shorter 2017; Mackenzie et al., 2017). Therefore, characterizing the role of LC domains in defining RBP structural states and assemblage formation in neurons is an important step towards elucidating the mechanisms that control proper local translation as well as RBP-mediated pathogenesis in the nervous system.

FXR2P (Fragile X related protein 2) is a member of the Fragile X related family of RNA binding proteins that also includes FXR1P and FMRP (Fragile X mental retardation protein). Variants in all three contribute to autism risk, with loss of FMRP causing the autism-related disorder Fragile X syndrome (Schluth-Bolard et al., 2010; Stepniak et al., 2015). All three FXR proteins have equivalent RNA binding properties via the KH domains (Darnell et al., 2009). However, FXR2P is unique in being an essential component of Fragile X granules (FXGs), a class of endogenous RNA granules that associate with ribosomes as well as mRNAs that encode proteins important for neuronal plasticity (Fragile X granules; Christie et al., 2009; Akins et al., 2017; Chyung et al., 2018). FXGs are only present in a subset of stereotyped neurons where they are restricted to axonal arbors (Akins et al., 2012; Akins et al., 2017).

These observations suggest that FXR2P harbors intrinsic features that contribute to its function in regulating higher order structure formation in neurons. The FXR2P amino terminal region is >90% identical to FMRP and contains two KH and Tudor RNA binding domains as well as nuclear import and export domains (Zhang et al., 1995; Darnell et al., 2009; Adams-Cioaba et al., 2010). However, FXR2P diverges from the rest of the Fragile X family in key respects as, for example, the FXR2P carboxyl-terminal domain has been reported to contain nucleolar-targeting signals (Tamanini et al., 1999; Tamanini et al., 2000). In our efforts to identify FXR2P domains critical for the formation of axonal RNA granules, we previously showed that FXR2P is the sole family member that is N-myristoylated, a modification that regulates its axonal distribution but not granule assembly (Stackpole et al., 2014). An intriguing possibility to regulate granule assembly is the carboxy-terminal low complexity domain that is a feature of all members of the Fragile X protein family (Kato et al., 2012). Notably, the FXR2P LC domain is divergent, sharing only ~19 and ~37% sequence identity respectively with the comparable regions of FMRP or FXR1P.

Here we sought to define the features of FXR2P that contribute to its role in mediating assemblage formation in neurons. We used an unbiased EGFP insertional mutagenesis strategy to delineate three discrete sites within a 23 amino acid region of the LC domain that promote the assembly of multiple distinct fibril types in neurons. These structures exist in both isolated and bundled forms and exhibit different developmental timelines, ultrastructure and state-dependent ribosome-RNA interactions. We find that FXR2P fibril formation within neurons requires elements within the LC domain as well as a functional KH2 RNA binding domain. Further, the mRNA and ribosome association varies between fibril type. We conclude that discrete elements within the FXR2P LC domain can mediate the formation of alternative higher-order structural states with differential ribosome association in neurons.

## RESULTS

### An unbiased insertional mutagenesis screen to identify FXR2P domains mediating assembly into higher order structural states

We performed an EGFP insertional mutagenesis screen to reveal FXR2P states important for mediating the formation of higher order assemblages in neurons. We anticipated that this approach could yield at least two broad classes of FXR2P^EGFP^ fusions that might: 1) mimic the typical granule structure observed endogenously; or 2) promote or disrupt the formation of higher order states. A library of FXR2P^EGFP^ insertional constructs was created using an *in vitro* transposition reaction that capitalizes on the random insertion of transposons to introduce EGFP coding sequence (CDS) into single sites within FXR2P (Fig. 1A; Sheridan et al., 2002; Giraldez et al., 2005). The library was screened for FXR2P^EGFP^ fusions containing in-frame, properly oriented EGFP at unique positions along the length of FXR2P CDS. This procedure yielded 18 different fusions that we term ‘FXR2P^[X]’^ where X is the amino acid position at which EGFP is inserted (Table 1; Fig. 1A).

**Table 1.** *Characterization of FXR2P^EGFP^ Constructs in Neurons*

| *Insertion Site* | *Domain* | *Diffuse Soma* | *Granule* | *Fibril* |
| --- | --- | --- | --- | --- |
| 15 | Tud1 | + | + | - |
| 20 | Tud1 | + | + | - |
| 26 | Tud1 | + | + | - |
| 36 | Tud1 | + | + | - |
| 65 | Tud2 | + | + | - |
| 77 | Tud2 | + | + | - |
| 88 | Tud2 | + | + | - |
| 94 | Tud2 | + | + | - |
| 127 | NLS | + | + | - |
| 158 | * | + | + | - |
| 217 | * | + | + | - |
| 218 | * | + | + | - |
| 341 | KH2 | + | + | - |
| 370 | NES | + | + | - |
| 416 | LC | - | - | + |
| 435 | LC | - | + | + |
| 439 | LC | - | + | + |
| 456 | LC | - | + | - |

*+, presence, -, absence. Abbreviations: Tud, Tudor domain; NLS, nuclear localization signal; \* uncharacterized; KH2, RNA binding KH2 domain; NES, nuclear export sequence; LC, low complexity*

**Figure 1.**
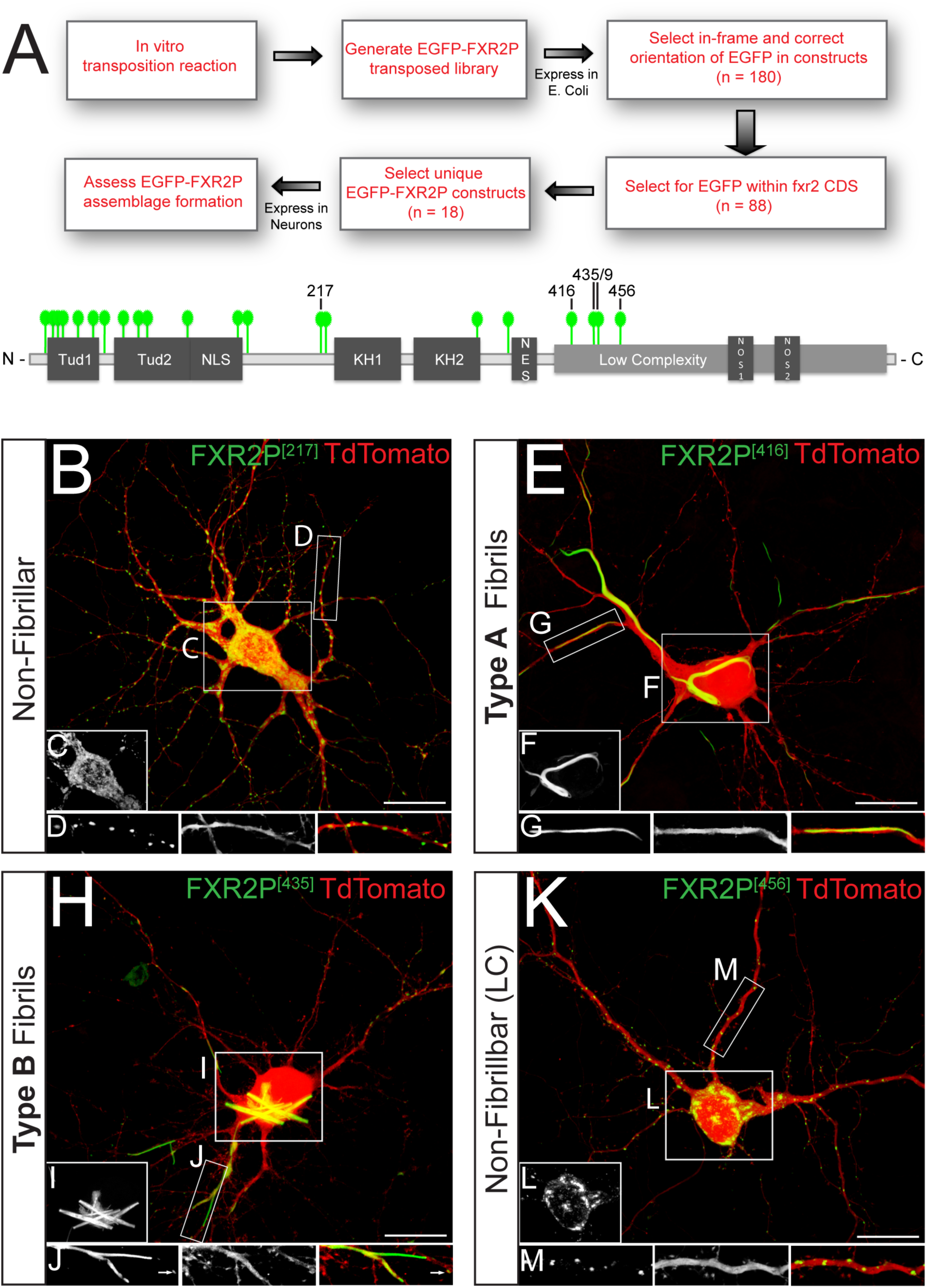
Unbiased insertional mutagenesis screen identifies a discrete region of the FXR2P LC domain as a regulator of assemblage formation in neurons. *(A)* Schematic of EGFP insertional mutagenesis approach used to generate a set of 18 unique, full-length EGFP-transposed FXR2P constructs (FXR2P^EGFP^; see Methods). Each green tag marks one specific EGFP insertion site within a FXR2P^EGFP^ construct. EGFP insertion at a given residue is denoted as FXR2P^[#]^ where ‘#’ is the amino acid insertion position in FXR2P. *(B-D)* Cultured cortical neuron co-transfected with FXR2P^[217]^ (green) and diffusible cell fill TdTomato (red). FXR2P^[217]^ is present diffusely in the soma (*inset C*) and in granules in dendrites (*inset D*). *(E-G)* FXR2P^[416]^ is present in Type A fibril bundles in both soma *(inset F)* and dendritic processes *(inset G)*. *(H-J)* FXR2P^[435]^ is present in dendritic granules (arrows) as well as Type B fibril bundles within soma *(inset I)* and dendritic processes *(inset J)*. *(K-M)* FXR2P^[456]^ is present in dendritic granules; no fibrils were observed in either soma *(inset L)* or processes *(inset M)*. All cultures DIV14. Scale bar = 20µm.

### EGFP fusions in the amino terminal half of FXR2P form granules localized to neuronal processes

To determine which domains of FXR2P influence its formation into higher order assemblages we assessed the cellular distribution of the 18 FXR2P^EGFP^ fusions expressed in cultured neurons (Fig. 1A; Table 1). Primary cortical neuron cultures were co-transfected at DIV3 (days *in vitro*) with each of the FXR2P constructs along with TdTomato to provide a diffusible cell fill. As summarized in Table 1, fusions with EGFP inserted into any of 14 different sites in the N-terminal half of FXR2P were diffusely distributed in cell somata and localized to discrete granules within dendrites and axons of DIV6 and DIV14 neurons (Fig.1B-D). This pattern was observed with EGFP fused into one of several regions within the N-terminal half of the molecule including the Tudor, RNA-binding KH2 and nuclear localization signal domains (Fig. 1A; Table 1). FXR2P^[217]^ was chosen as the representative of this set of 14 N-terminal fusions (Fig. 1B-D). The cellular distribution of FXR2P^[217]^ is qualitatively similar to that previously observed for heterologously-expressed FXR2P (Levenga et al., 2009; see also Stackpole et al., 2014). Taken together, these data indicate that FXR2P with EGFP inserted within N-terminal domains organizes into granular assemblages within neurons.

### EGFP insertion within a restricted region of the LC domain results in the formation of two distinct fibrillar states of FXR2P

FXR2P with EGFP fused within a restricted region of the C-terminal LC domain assumed strikingly different organizations when expressed in neurons. When viewed at the light microscopic level, FXR2P fusions at residue 416 (FXR2P^[416]^) assembled into elongated, curvilinear structures that we term Type A fibril bundles (Fig. 1E-G; see ultrastructural analysis below). Type A fibril bundles were thread-like, apparently flexible structures present in somata, axons and dendrites. Neurons containing Type A fibril bundles exhibited few to no FXR2P^[416]^ granules within processes. Moreover, little diffuse signal was observed anywhere in the cell, suggesting that the large majority of FXR2P^[416]^ had assembled into the fibril bundles. FXR2P^[416]^-expressing neurons exhibited Type A fibril bundles at all times examined (DIV6-28), with their length and complexity increasing with age (Fig. 2A-B). Together, these data demonstrate that FXR2P^[416]^ adopts a distinctive fibrillar organization across multiple developmental stages in cultured neurons.

**Figure 2.**
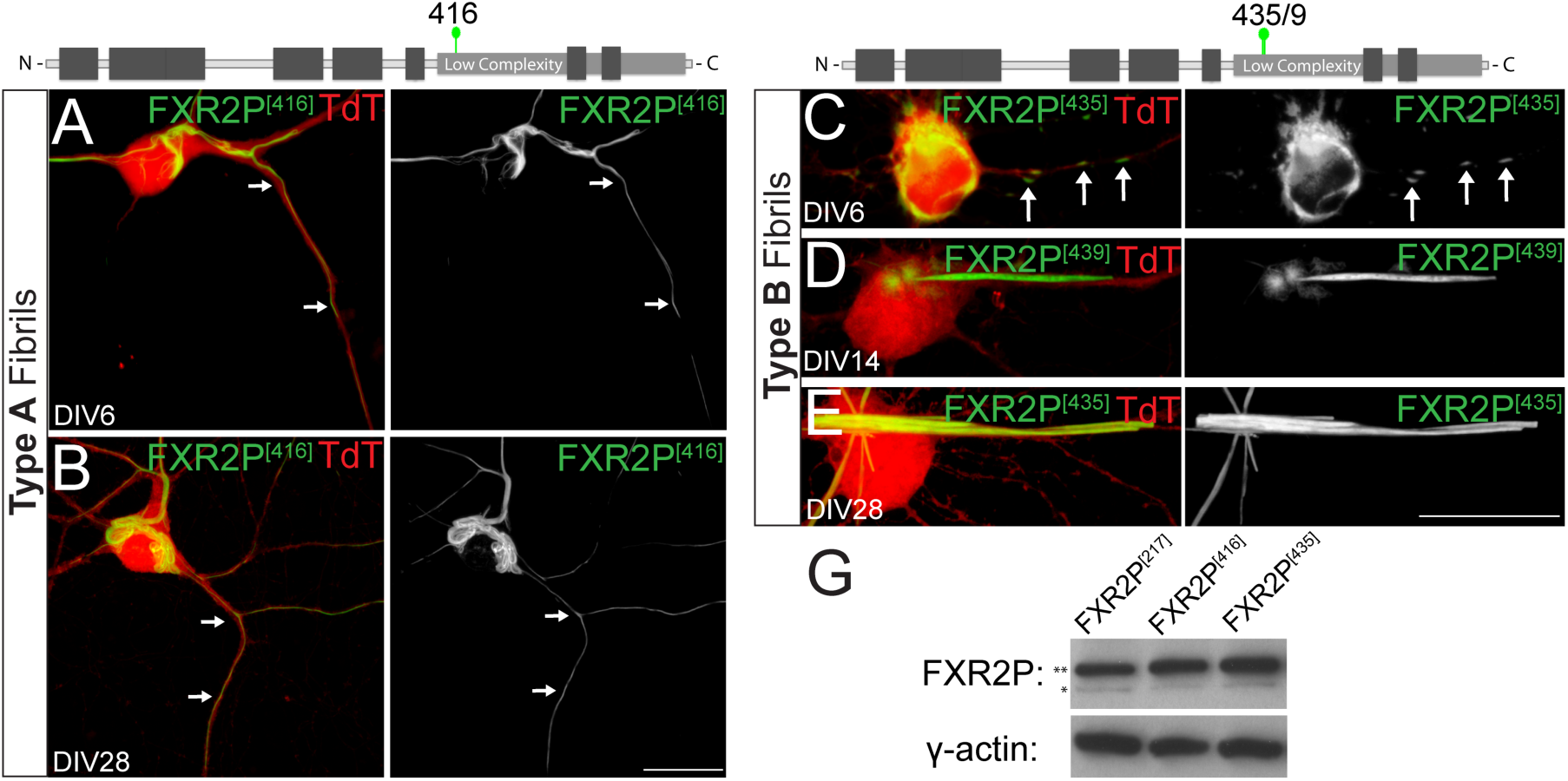
Time course of Type A and B fibril bundle expression in cultured neurons. *(A)* DIV6 neuron co-transfected with FXR2P^[416]^ (green) and TdTomato (red). Type A fibril bundles are detected in the cell body and extending into dendritic processes. Arrows mark localization of fibrils in dendrites. Dendritic FXR2P granules are not observed. *(B)* Type A fibril bundles observed in soma and dendrites of a DIV28 neuron. Note that the length of fibrils increases both in the cell soma and processes compared to DIV6. Arrows mark FXR2P fibrils in a dendrite. *(C)* Dendrite of a DIV6 neuron co-expressing FXR2P^[435]^ (green) and TdTomato (red). FXR2P^[435]^ is present in dendrites only in granules at this time (arrows). *(D)* In DIV12 neurons, FXR2P^[435]^ localizes to discrete, spherical, nest-like structures with a fibrillar substructure. These nest-like structures are closely associated to newly-forming Type B fibril bundles. *(E)* FXR2P^[435]^ forms Type B fibril bundles in a DIV28 neuron that extend from the cell body into dendritic processes. *(G)* Western blot analysis of protein lysates from DIV6 neuronal cultures expressing FXR2P^[217]^, FXR2P^[416]^ or FXR2P^[435]^ probed with antibodies to FXR2P and GAPDH (loading control; 37kD). Note that all the FXR2P^[EGFP]^ fusions are expressed at similar levels. Equivalent results were observed in two independent experiments. The anti-FXR2P detected both the FXR2P^[EGFP]^ fusions (upper band ~122kD; double asterisk) as well as endogenous FXR2P (lower band ~95kD; single asterisk). Scale bar = 20µm.

EGFP insertions at either residue 435 or 439 in the LC domain assembled into a second remarkable structure when expressed in neurons that we term Type B fibril bundles (Fig. 1H-J; Fig. 2C-E; and see below). Since fusions at either of these two positions yielded similar results, we will use the term ‘FXR2P^[435/439]’^ to describe their properties; specific constructs used for given experiments are noted in the Figures. Type B fibril bundles were straight and crystal-like structures. These bundles were observed at ≥DIV11 and were present in both somata and dendrites (Fig. 2C-E). Type B fibril bundles were observed for up to four weeks in culture with bundle size increasing over time (Fig. 2C-E). FXR2P^[435/439]^ was also localized to smaller dendritic granules at all time points investigated (Figs 1 and 2). FXR2P^[435/439]^ was also present within spherical, nest-like, finely fibrous structures that were closely associated with Type B fibril bundles (Fig. 2D). These nest-like structures, reminiscent of dandelion seed pods, were only observed from ~DIV9-14. Finally, FXR2P fusions at residue 456 within the LC domain (FXR2P^[456]^) formed neither fibrillar nor nest-like structures in neurons. Rather, FXR2P^[456]^ was localized to granules that were widely distributed within the cell somata and neuronal processes (Fig. 1K-M), similar to that observed with FXR2P^[217]^. Thus, EGFP insertion into a discrete region within the LC domain of FXR2P (residues 416-439) resulted in formation of two novel types of fibrillar structures in neurons.

We considered the possibility that the fibrillar bundles formed by FXR2P^[416]^ and FXR2P^[435/439]^ might be caused by differential expression of these fusions and/or reflect reduced cell viability. However, western blotting demonstrated that all the fusions tested showed equivalent expression levels (Fig. 2G). Moreover, this biochemical analysis indicated that all FXR2P^EGFP^ fusions tested were intact, with no signs of cleavage. Further, as judged by the TdTomato cell fill, neurons expressing any of the FXR2P^EGFP^ fusions demonstrated comparable somatodendritic and axonal morphologies (Fig. 1) for up to four weeks in culture (Fig. 2). Taken together, these observations suggest that Type A and B fibril bundles most likely form due to intrinsic differences in protein assembly caused by EGFP insertion into the LC domain.

### FXR2P is an intrinsically disordered protein with a non-prion-like LC domain

The N-terminal half of FXR2P shares high sequence similarity and domain organization with FMRP, an extensively characterized protein (Zhang et al., 1995; Darnell et al., 2009; Adams-Cioaba et al., 2010). In contrast, little is known about the C-terminal half of FXR2P. We therefore analyzed FXR2P using the bioinformatic predictors of intrinsically unfolded and disordered regions PONDR-FIT and FoldIndex (Xue et al., 2010; Prilusky et al., 2005). Both algorithms predicted that the C-terminal half of FXR2P harbors an LC domain with an intrinsically disordered and unfolded structure (residues ~388-673; Fig. 3A, D). We next determined whether the FXR2P LC domain was predicted to harbor the intrinsic ability to fibrillize into steric zipper structures. ZipperDB (Goldschmidt et al., 2010) predicted multiple residues in the LC domain with a high propensity for fibrillization (Fig. 3B). Interestingly, one of these regions, residues 415-419 (ESSSS; Fig. 3B) was coincident with the EGFP insertion site that results in Type A fibril formation (FXR2P^[416]^, see above). A high contribution of residues prone to pi-pi contact formation (Fig. 3C) suggests that FXR2P may self-interact like other pi-rich proteins that undergo phase separation, a critical regulatory feature of LC-driven self-assembly and organization of higher order macromolecules (Mitrea and Kriwacki 2016; Chong and Forman-Kay 2016; Vernon et al., 2018). Taken together with the ZipperDB analyses, the LC domain of FXR2P is thus predicted to be primed for phase separation and/or intrinsic self-assembly.

**Figure 3.**
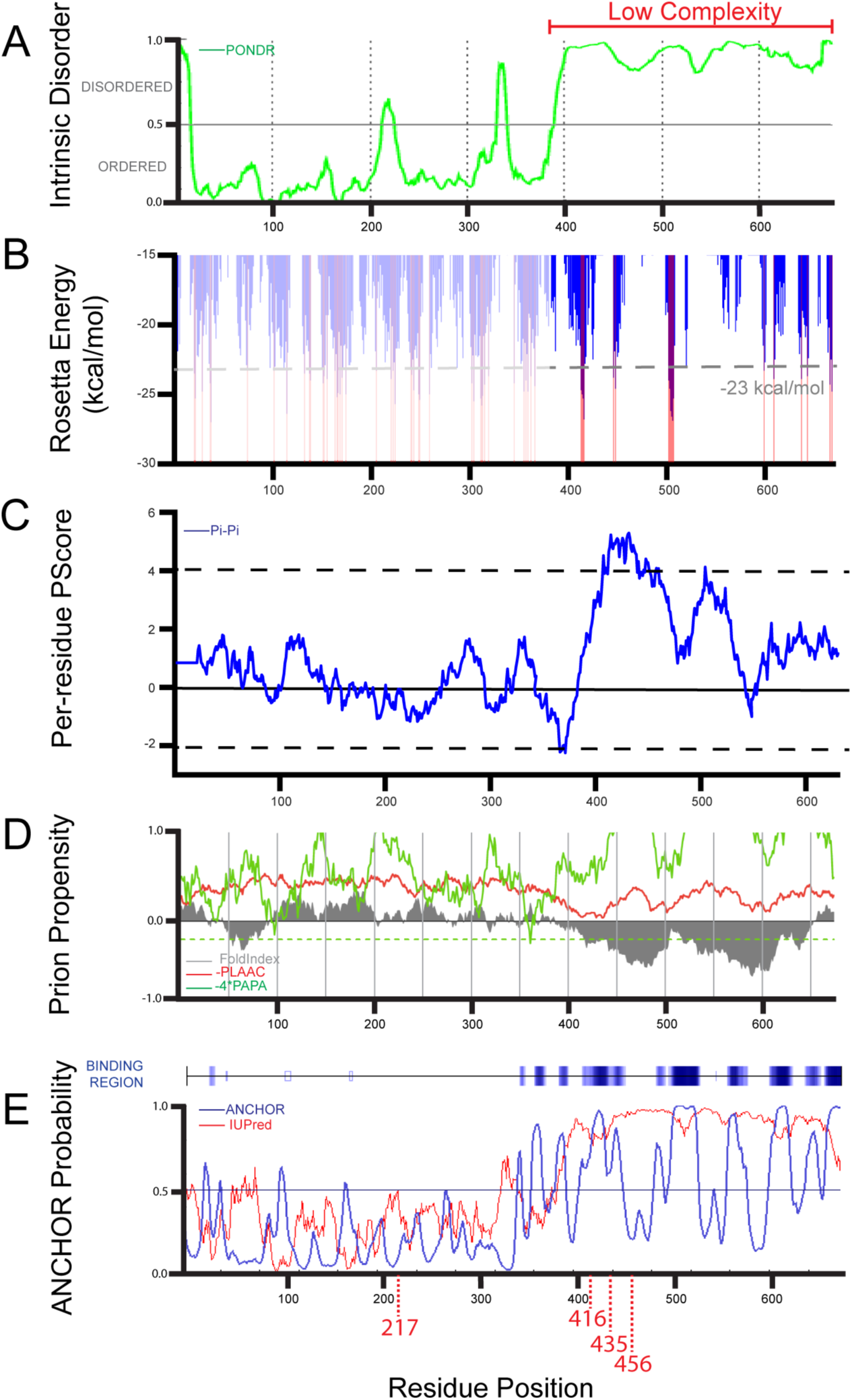
FXR2P contains a C-terminal LC domain that is intrinsically disordered. *(A)* PONDR-FIT predicts that a low complexity region in the C-terminus of FXR2P (residues 388-673) is intrinsically disordered. PONDR-FIT scores > 0.5 indicate a probability towards intrinsic disorder. (*B*) ZipperDB predicts that regions around residues 415 and 500 have increased fibrillization propensity. (Note that the apparent increased fibril forming propensity of regions N-terminal to the low complexity domain [light shading] are due to the folded nature of these domains and are not relevant to this analysis.) (*C*) Phase separation prediction based on the per-residue pi-contact propensity indicates that the low complexity region of FXR2P has increased propensity to form pi-contacts compared to its folded domains. Dotted lines represent PScore thresholds for enrichment of pi-contacts (PScore ≥ 4) or depletion of pi-contacts (PScore ≤ −2). *(D)* FoldIndex predicts that the LC region of FXR2P is intrinsically unfolded (grey shading less than zero). However, no region within FXR2P was predicted to be prion-like by either the PLAAC algorithm (red; log-likelihood ratio score below zero) or the PAPA algorithm (green; log-odds ratio score below dashed green line). *(E)* ANCHOR predicts multiple disordered binding regions within the LC domain of FXR2P (blue; score > 0.5; darker blue signifies higher ANCHOR score). IUPRED predicts the LC domain as intrinsically disordered segment (red; score > 0.5 are predicted as disordered). Arrows below the residue position denote the location of the 217, 416, 435/439 and 456 EGFP insertion sites.

We also asked whether the FXR2P LC domain contained prion-like elements, which are characteristic of RBPs implicated in neurodegenerative disease (Harrison & Shorter 2017; King, Gitler & Shorter 2012). However, Fig. 3D shows that the FXR2P LC domain lacks Q/N rich regions as judged by either the PAPA or PLAAC algorithms (Toombs et al., 2012; Lancaster et al., 2014). Finally, we used ANCHOR to predict the locations of disordered binding regions within the FXR2P LC domain (Dosztanyi et al., 2009). This algorithm failed to detect favorable intrachain interactions in the LC domain that might promote folding into a well-defined structure, but did identify disordered protein segments that could undergo a disorder-to-order transition upon binding with globular protein partners (Fig. 3E; Meszaros et al., 2009). Notably, such predicted *trans* domains were present within the discrete region targeted in the fusion constructs (positions 416 – 439), but were absent from the position 456 region (Fig. 3E). Together, these algorithms indicate that FXR2P is an intrinsically disordered protein containing a large, non-prion-like LC region constituting ~42% of its length and has the potential to restructure into defined three-dimensional shapes upon protein binding.

### FXR2P LC fusions form distinct fibril types that assemble into bundles with differential ribosome association

We next investigated the ultrastructure of neurons expressing FXR2P^[217]^, FXR2P^[416]^ or FXR2P^[435]^ (granules only, Type A fibrils, or Type B fibrils, respectively; see above). Neurons expressing FXR2P^[217]^ contained discrete structures within dendrites that were rich in ribosomes embedded in electron-dense, amorphous material (Fig. 4D). These structures are likely to correspond to the granules observed by light microscopy (Fig. 1). In addition, neuronal somata contained numerous polysomes (Fig. 4A). No fibrillar structures were observed in cells expressing FXR2P^[217]^.

**Figure 4.**
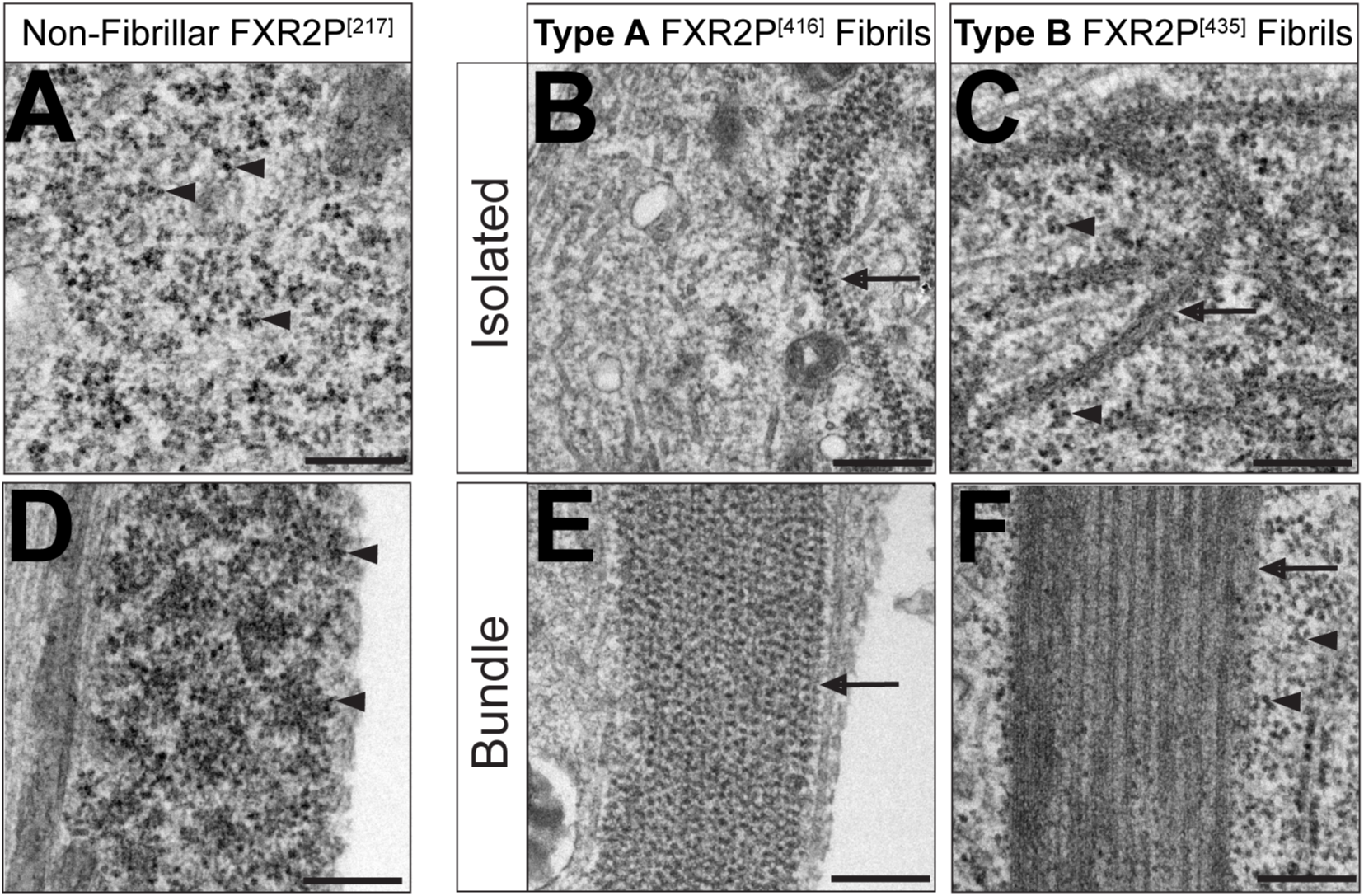
Ultrastructure of FXR2P^[EGFP]^ Type A and B isolated fibrils and fibril bundles in neurons. Electron microscopy of neurons transfected with non-fibrillar FXR2P^[217]^ *(A, D)*; Type A fibril-forming FXR2P^[416]^ *(B, E);* or Type B fibril-forming FXR2P^[435]^ constructs *(C, F). (A)* Cytoplasm of DIV6 neuron expressing FXR2P^[217]^ is rich in polysomes (arrowheads) and no fibrils are observed. *(B)* Isolated Type A fibrils decorated with ribosomes (arrows) from DIV6 neuron transfected with FXR2P^[416]^. Note that polyribosomes are not observed in the cytoplasm. *(C)* DIV14 neuron transfected with FXR2P^[435]^ displays ultrastructurally distinct, isolated Type B fibrils decorated with ribosomes (arrows). Polyribosomes are observed in the cytoplasm (arrowhead). *(D)* A dendritic granule in an neuron expressing FXR2P^[217]^ enriched with ribosomes (arrowhead). *(E)* Type A fibril bundle (arrow) associated with ribosomes in a neuron expressing FXR2P^[416]^. *(F)* Type B fibril bundle (arrow) devoid of ribosomes in a neuron expressing FXR2P^[435]^. Note that polysomes are readily observed in the adjacent cytosol (arrowheads). Scale bar = 250nm.

Electron microscopy of neurons expressing FXR2P^[416]^ revealed distinctive cytoplasmic fibrils that were present in both isolated and bundled configurations (Fig. 4B, 4E; Fig. 5A-C). Isolated Type A fibrils were strand-like and thin (width: 20.5 ± 0.7 nm; mean ± SEM; n = 9 fibrils, 3 neurons). These strand-like structures were observed coursing within the cytoplasm of cell bodies (Fig. 4B) and were frequently adjacent to perinuclear amorphous, electron dense material (Fig. 5A-B). This co-localization suggests that this electron dense material could act as a zone for fibril production.

**Figure 5.**
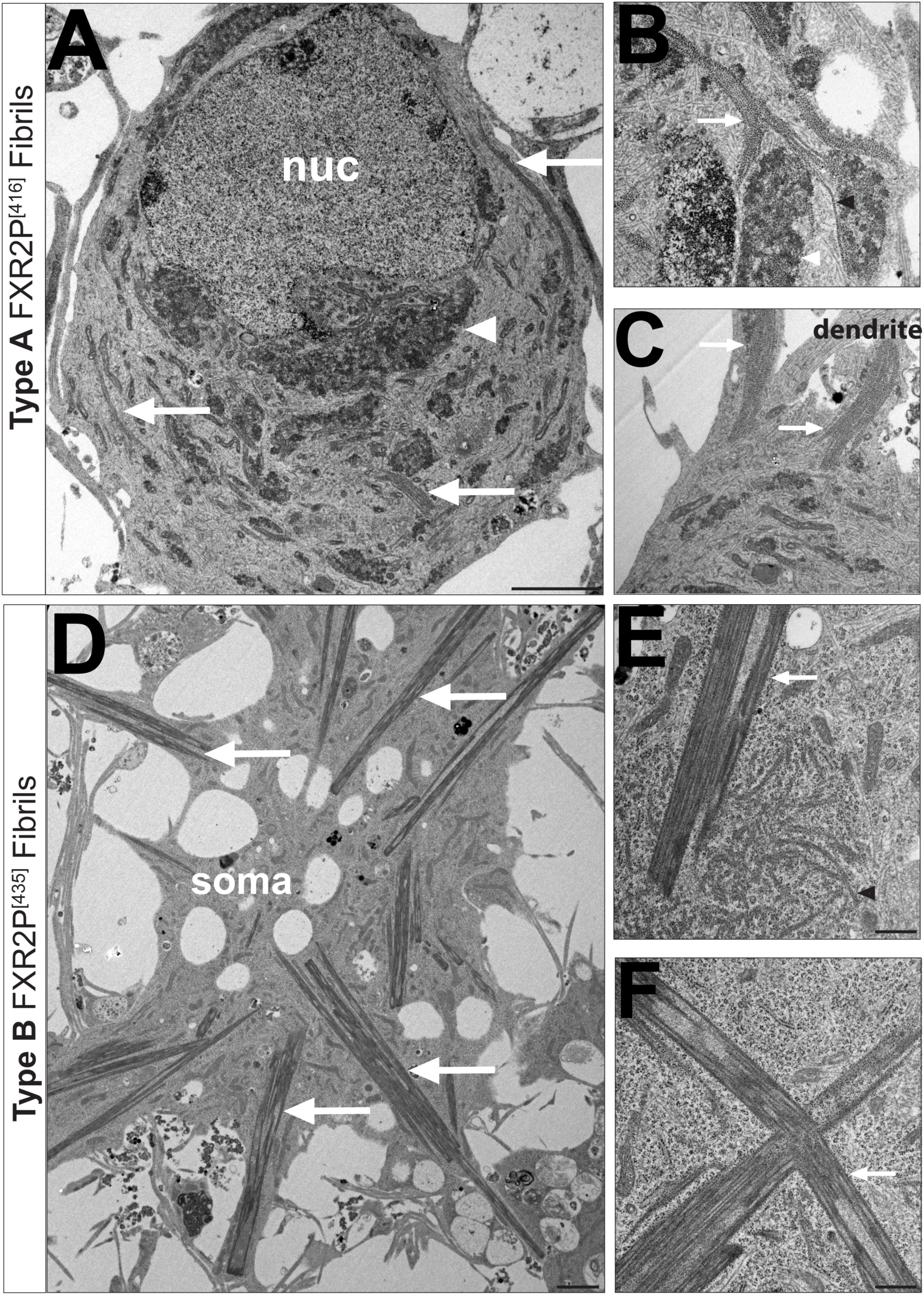
Disposition and ultrastructural localization of Type A and B FXR2P isolated fibrils and fibril bundles in neurons. *(A-C)* Electron micrographs of DIV6 cortical neurons transfected with Type A fibril-forming FXR2P^[416]^. *(A)* Low magnification view of cell soma shows Type A fibril bundles coursing through cytoplasm (arrows). Juxtanuclear amorphous material is also observed (white arrowhead). *(B)* Isolated Type A fibrils (black arrowhead) and bundles (arrows) in close proximity to granular material (white arrowheads). Note the apposition of the isolated Type A fibril with the granular domain. *(C)* Type A fibril bundles (arrows) extend from cell soma into dendritic processes. *(D-F)* Electron micrograph of DIV14 cortical neurons expressing Type B fibril-forming FXR2P^[435]^. *(D)* Low magnification view of cell shows multiple rigid Type B fibrils extending radially from cell center (arrows). Plane of section is adjacent to the substrate. *(E)* Isolated Type B ribosome-decorated fibrils (black arrowhead) are restricted to a circular domain in the cytoplasm, which is likely to be a section of the nest-like structures observed by fluorescence microscopy (Fig. 2D). Note that a ribosome-free Type B fibril bundle is juxtaposed to this nest-like structure. *(F)* Type B fibril bundles are rectilinear and jagged in the cytoplasm as well as devoid of ribosomes. Scale bar in *A, B* = 2µm; scale bar in *B, C, E, F* = 500nm.

Type A fibrils were also present in highly organized bundles (Fig. 4E; Fig. 5B-C). These fibril bundles were wide structures (Fig. 4E) observed throughout the somata and extending into dendrites (Fig. 5A-C). The size and disposition of these fibril bundles indicate that they correspond to the thread-like structures observed by fluorescence microscopy (Figs. 1 and 2). Taken together, these observations indicate that isolated Type A fibrils assemble into bundles with no gross changes in their basic structure.

Ultrastructural analysis of neurons expressing Type B fibril-forming FXR2P^[435]^ revealed a strikingly different class of isolated fibrils and fibril bundles. Isolated Type B fibrils were short and arc-like (average width: 41.6 ± 1.5 nm; n = 12 fibrils, 2 neurons; Fig. 4C) and clustered within discrete spherical zones (Fig. 5E). These zones are likely to correspond to the spherical, nest-like structures observed to be adjacent to Type B fibrils bundles by fluorescence microscopy (Fig. 2D).

Neurons expressing FXR2P^[435]^ also exhibited highly regular fibril bundles with an overall needle-like appearance as they coursed through the cytoplasm (Fig. 4F; Fig. 5D-F). Type B bundles extended radially from cell somata into processes (Fig. 5D) and were comprised of tightly aligned individual fibrils (Fig. 4F). This observation suggests that isolated Type B fibrils can transition from a short, arc-like state to a rigid, elongated configuration when assembled into the Type B bundles. Finally, the clusters of isolated Type B fibrils were often juxtaposed to fibril bundles (Fig. 5E), suggesting that these zones may be organizing centers for fibril bundling.

### Type A and B fibril bundles show differential ribosome association

We next assessed the ribosomal association of the FXR2P fibrils. Figure 4 shows that ribosomes associated with both isolated and bundled Type A fibrils (FXR2P^[416]^) in a periodic fashion with a spacing of 28.9 ± 1.5 nm and 31.4 ± 0.7 nm, respectively (n = 10 and 14 fibrils, respectively, 3 neurons; Fig. 4B, 4E). Remarkably, electron microscopy also revealed the wholesale redistribution of ribosomes within cells expressing FXR2P^[416]^. In contrast to neurons expressing FXR2P^[217]^, where polysomes were widely distributed in the cell soma (Fig. 4A), neurons expressing FXR2P^[416]^ showed ribosomes decorating Type A fibrils and were not observed free in the cytosol (Fig. 4B, 4E).

Isolated and bundled Type B fibrils showed strikingly differential ribosome association. Although ribosomes associated with isolated Type B fibrils (Fig. 4C), Type B fibril bundles were devoid of ribosomes (Fig. 4F). Further, ribosomes were abundant in the cytosol of neurons containing Type B fibrils (Fig. 4F; Fig.5E-F). Taken together, these results show that manipulation of specific sites within the FXR2P LC domain results in the formation of two structurally distinct, ribosome-decorated isolated fibril types (A and B). However, when assembled into bundles, FXR2P fibrils either associate with (Type A) or are devoid of (Type B) ribosomes. Further, virtually all ribosomes in neurons expressing FXR2P^[416]^ partition to Type A fibrils.

We used fluorescent microscopy to confirm and extend the ultrastructural observations of ribosome association with FXR2P fibrils. We first asked whether FXR2P^[217]^ colocalized with either rRNA or polyA^+^ RNA in neurons. Using a monoclonal antibody that recognizes 5S/5.8S rRNA (Y10b; see Methods), we observed that a subset of FXR2P^[217]^ colocalized with rRNA within dendritic granules of DIV14 neurons (Fig. 6A). Further, labeling with an oligo(dT) probe demonstrated that a subset of dendritic FXR2P^[217]^ granules colocalized with polyA^+^ RNA (Fig. 6B). Colocalization was also observed for Type A fibril bundles with 5S/5.8S rRNA and polyA^+^ RNA (Fig. 6C-D). No signal was observed when sense probes were used (Fig. 6E).

**Figure 6.**
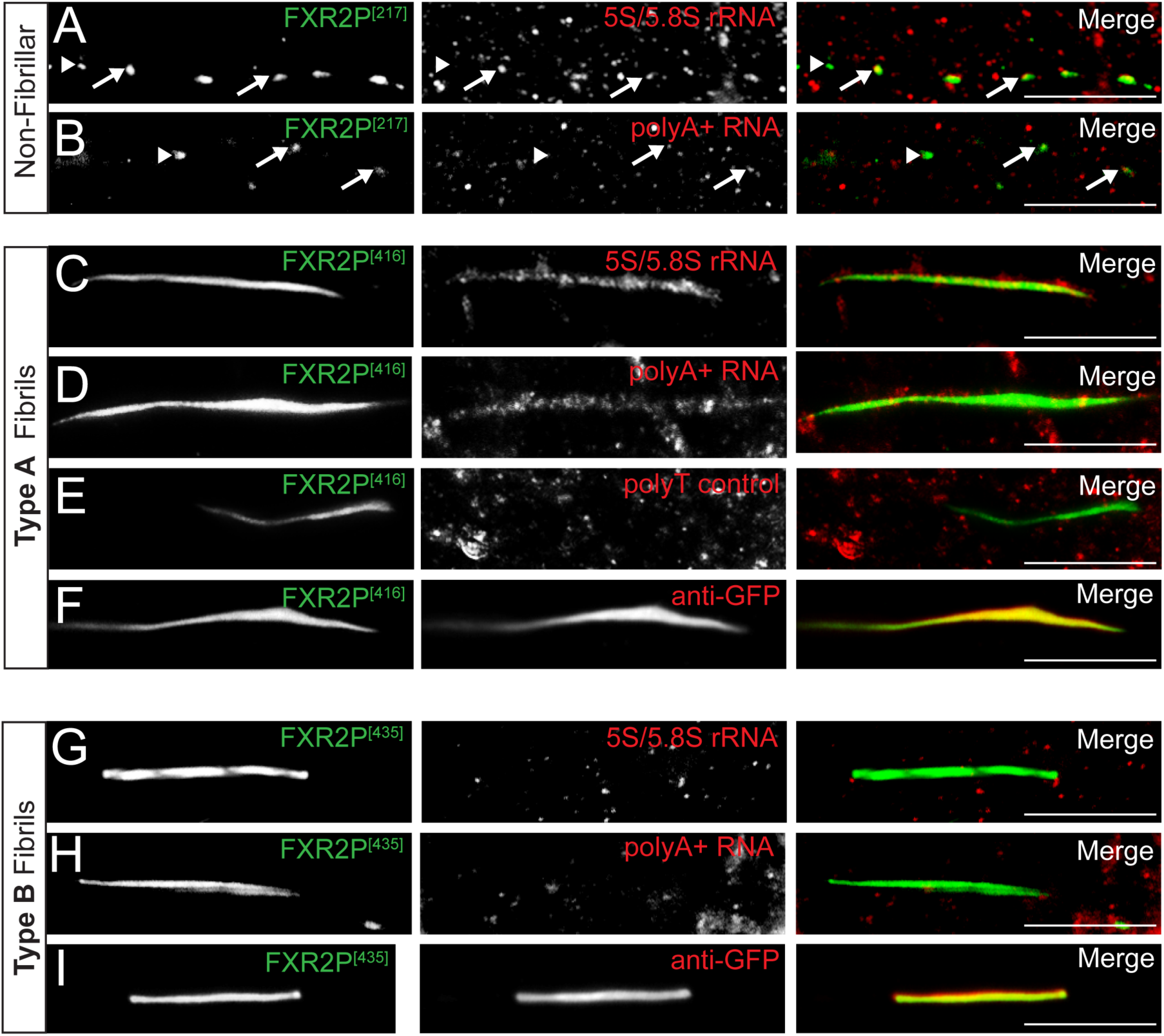
Ribosomes and mRNA co-localize with Type A but not Type B fibril bundles. *(A, B)* FXR2P^[217]^ granules colocalize with 5S/5.8S rRNA (Y10b antibody; red; *A*) and polyA^+^ mRNA (red; *B*) in dendrites. *(C-E)* Type A fibrils colocalize with 5S/5.8S rRNA (red; *C*) and polyA+ mRNA (red; *D*) in dendrites. No signal was observed in fibrils with a polyT control (red; *E*). *(G, H)* Type B fibrils do not colocalize with either 5S/5.8S rRNA (red; *G)* or polyA^+^ mRNA (red; *H*). *(F, I)* Anti-GFP immunostaining (red) of Type A (*F*) and Type B (*I*) fibril bundles. Note the complete co-localization of both the FXR2P^[416]^ and the FXR2P^[435]^ intrinsic EGFP signals (green) with the anti-GFP immunostain (red). DIV14 neurons. Scale bar = 10µm.

Finally, in agreement with our ultrastructural analyses no rRNA or polyA^+^ RNA was detected in Type B fibril bundles (Fig. 6G-H). We considered the possibility that the rigid and tightly ordered structure of Type B fibrils might restrict antibody accessibility. To address this potential confound, we performed immunofluorescence with an anti-GFP antibody and observed the signal from this reagent was indistinguishable from that of the respective FXR2P^EGFP^ fusions (Fig. 6F, 6I). These findings provide further evidence that manipulation of the FXR2P LC domain results in the assembly of two distinct higher order states in neurons distinguishable by their structure and ability to associate with RNA.

### Both RNA binding and LC domains are required for FXR2P fibril formation in neurons

We next asked whether the FXR2P LC domain is necessary for assemblage formation in neurons. As shown in Figure 7, a mutant FXR2P lacking the LC domain (FXR2P^[217∆LC]^) was distributed diffusely throughout the cell somata and processes. Neither granules nor fibrils were observed (Fig. 7A-C). Combined with the above results, these data indicate that the FXR2P LC domain is required for its ability to form higher order assemblages within neurons.

**Figure 7.**
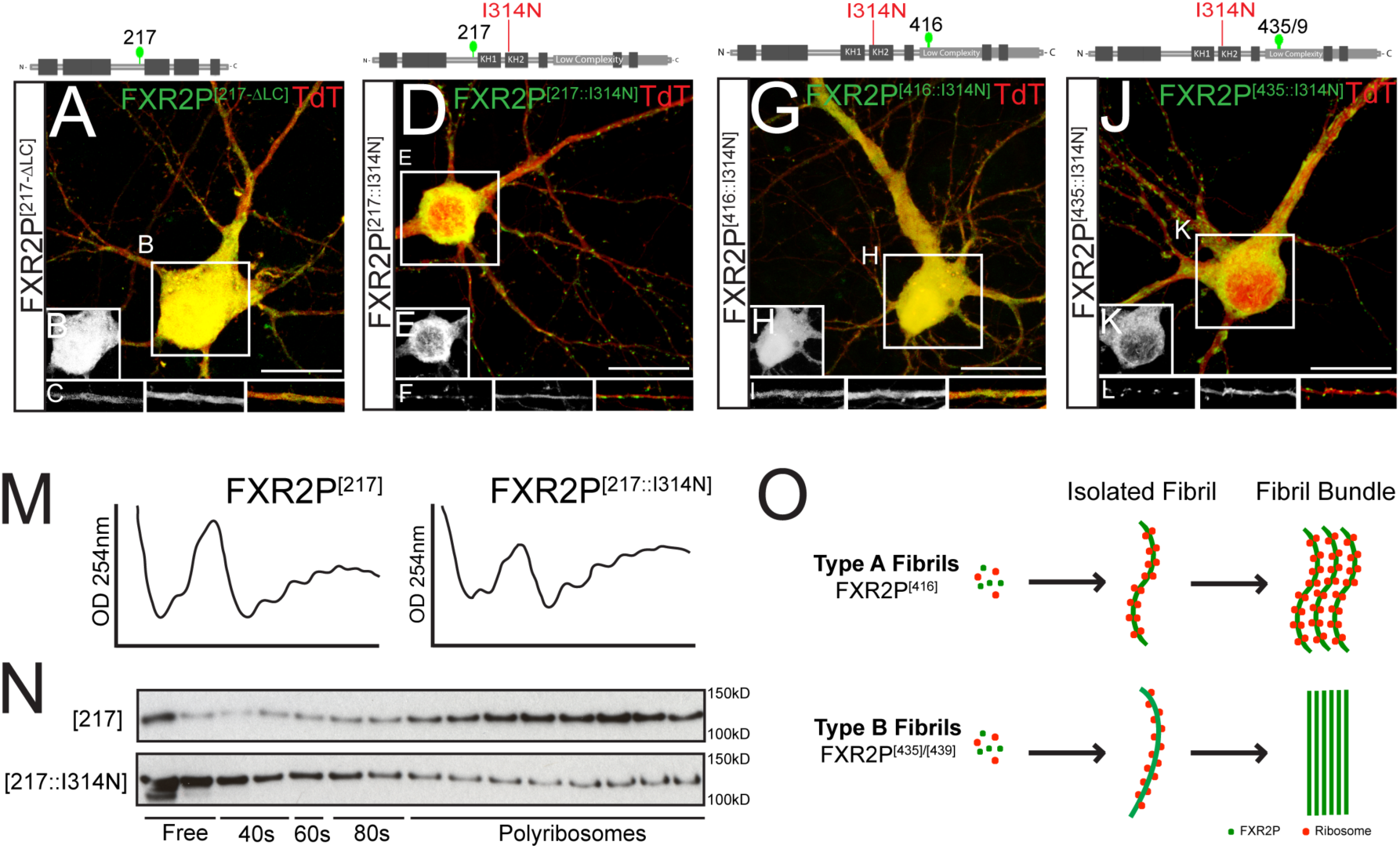
FXR2P RNA binding and LC domains collaborate in fibril formation. *(A-* DIV14 neuron co-transfected with (FXR2P^[217∆LC]^; green) and TdTomato (red). FXR2P^[217∆LC]^ is diffusely localized throughout the cell and discrete granules are not observed (*insets B* and *C, respectively*). *(D-F)* FXR2P^[217::I314N]^ is diffusely localized in the soma *(D; inset E)* and present in granules in cell processes *(inset F).* Compare to Fig. 1B-D. *(G-I)* FXR2P with a point mutation in the RNA-binding KH2 domain (FXR2P^[416::I314N]^; green) is expressed diffusely in the nucleus and cell processes (insets *H* and *I,* respectively). No granular or fibrillar structures are observed. Compare to Fig. 1E-G. *(J-L)* FXR2P^[435::I314N]^ (green) is diffusely distributed in the soma (*J; inset K)* and is present in granules in dendrites *(inset L).* No fibrillar structures are observed. Compare to Fig. 1H-J. *(M)* A_254_ traces of sucrose gradients from HEK293T cells expressing either FXR2P^[217]^ or the RNA binding mutant FXR2P^[217::I314N]^*. (N)* Western blot for FXR2P in fractions collected from sucrose gradients from either FXR2P^[217]^ (upper blot) or FXR2P^[217::I314N]^ (lower blot). FXR2P^[217]^ predominantly co-sediments in polysome fractions. In contrast, FXR2P^[217::I314N]^ is enriched in free and monosome fractions and present in low levels in polysome fraction. Equivalent results were observed in two independent experiments. *(O)* Summary of Type A and B fibril dynamics and ribosome association in neurons. See text for details. Neurons transfected at DIV3 and analyzed on DIV14 neurons. Scale bar = 20µm.

We then sought to investigate the role of FXR2P RNA binding in assemblage formation. Mutations that abrogate FXR2P RNA binding have not been reported. However, the FXR2P KH2 domain is over 90% identical to that in FMRP (Zhang et al., 1995). Moreover, an I304N point mutation in the FMRP KH2 domain abrogates its RNA binding and ribosome association (Zhang et al., 1995; Feng et al., 1997, Laggerbauer et al., 2001; Darnell et al., 2005a; Darnell et al., 2005b; Zang et al., 2009; Ascano et al., 2012). We therefore mutated the comparable residue in FXR2P (I314N). To directly test whether this I314 mutation affects polysome association, we analyzed ribosome co-sedimentation by sucrose gradients from extracts of HEK293T cells expressing either FXR2P^[217]^ or FXR2P^[217::I314N]^. Figure 7M and 7N show that FXR2P^[217]^ was enriched in polysome fractions. This sedimentation profile is in agreement with previous work demonstrating that FXR2P predominantly co-fractionates with polysomes (Feng et al., 1997; Darnell et al., 2009; Zang et al., 2009). In contrast, FXR2P^[217::I314N]^ was shifted to lighter, non-polysome fractions (Fig. 7N). The A_254_ traces were similar between FXR2P^[217]^ and FXR2P^[217::I314N]^ extracts, indicating that global translation was similar in cells expressing either FXR2P form (Fig. 7M). Together, these results indicate that the I314N mutation perturbs FXR2P RNA binding.

We then assessed the role of FXR2P^EGFP^ RNA binding in neuronal assemblage formation. As shown in Figure 7D-F, FXR2P^[217::I314N]^ was distributed diffusely in cell bodies and was localized to granules in dendrites, a comparable localization to that observed with FXR2P^[217]^ (see Fig. 1). Thus an intact LC domain, but not RNA binding ability, is required for the formation of granular FXR2P assemblages in neurons. In contrast, the RNA-binding mutant FXR2P^[416::I314N]^ was diffusely distributed in the somata and cell processes; neither fibril bundles nor granular assemblages were observed (Fig. 7G-I). Finally, FXR2P^[435::I314N]^ was present in granular assemblages within the neuronal processes and was localized diffusely in the somata. No fibrillar structures were observed, even after DIV14 (Fig. 7J-L). These compact dendritic granules were similar to those observed in neurons expressing FXR2P^[435/9]^ at DIV6-28 (See Fig. 2C). Moreover, the fibrous, nest-like structures observed at >DIV9 were not detected (Fig. 7J-L). Taken together, these data indicate that collaboration between distinct regions within the LC and the RNA-binding domain are required to drive formation of both Type A and Type B fibril bundles in neurons.

## DISCUSSION

In this study we demonstrate that the RNA binding protein FXR2P can assume multiple higher-order states in neurons with unique ultrastructure, developmental timelines and ribosome/RNA association (Fig. 7O). Formation of these assemblages requires coordination between elements in the LC domain and a functional KH2 RNA binding domain. Our results highlight the important role of discrete LC regions in regulating the formation of RBP complexes in a neuronal context and suggest mechanisms by which differential structural and higher order states of an RBP can influence its protein and RNA association.

We utilized an insertional mutagenesis approach to identify the FXR2P LC domain as a key regulator of its assembly into complexes in neurons. Several observations indicate that the structures observed in our study reflect LC-dependent FXR2P states are not due to EGFP insertion per se. First, of the 18 EGFP fusions studied, 15 had no detectable fibrillar organization when expressed in neurons (Table 1). Instead, fibrillar FXR2P was observed in only 3 fusions that harbored insertions within a 23 amino acid stretch of the LC domain (residues 416-439; Figs 1 and 3). Second, an FXR2P fusion with EGFP inserted at position 456 in the LC domain (Fig. 1) did not assemble into fibrils. Third, as discussed below, FXR2P^[416]^ or FXR2P^[435]^ only formed fibrils when the KH2 RNA binding domain was functionally intact. Fourth, fusions that assemble into vastly different structural states were expressed at similar levels (Fig. 2). Fifth, deletion of the LC domain resulted in the complete loss of FXR2P assemblages in neurons (FXR2P^[217∆LC]^; Fig. 7). Finally, FXR2P fibrillization was not secondary to poor viability as neurons expressing either FXR2P^[416]^ or FXR2P^[435]^ were healthy for the 4 week observation period and showed highly elaborate dendritic and axonal arbors (Fig. 2).

This work identifies four novel FXR2P assemblages: Type A and Type B fibrils, which each exist both in isolation and in bundles (summarized in Fig. 7O). Ultrastructural analyses show that isolated A and B fibrils have distinct morphologies with unique shapes and diameters (~20nm and ~42nm, respectively). Type A bundles are thread-like and curvilinear structures that course within the cell body, dendrites and axons at all observed time points (DIV6-28). In contrast, Type B bundles are only found in ≥DIV11 neurons and are straight, needle-like structures that extend radially from cell somata into processes.

The FXR2P^EGFP^ fibrils described here are morphologically distinct from those observed in previous studies examining LC domain-containing RBPs. For example, RBPs such as FUS and hnRNPA1/2 form fibers that are ~30nm in diameter and shaped similarly to cross-β structures prototypical of amyloid fibers (Kato et al., 2012; Kim et al., 2013; Schwartz et al., 2013; Hughes et al., 2018; Luo et al., 2018). One possible explanation for the unique morphology of FXR2P^EGFP^ fibrils is a difference in the sequence of its LC domain. The FXR2P LC domain lacks repetitive [G/S]Y[G/S] motifs and QN-rich prion like sequences, both of which are linked to formation of amyloid-like fibrils (Hughes et al., 2018; Luo et al., 2018; Sun et al., 2011; Malinovska et al., 2013). Intriguingly, RBPs that form cross-β fibers are commonly implicated in aggregate pathology observed in neurodegenerative disorders (Ramaswami et al., 2013; Li et al., 2013; Aguzzi and Altmeyer 2016). However, neurons containing either FXR2P^EGFP^ fibril type survived at least 28 days in culture without significant cell death. Taken together, an interesting prospect is that the unique morphologies and non-pathogenicity of FXR2P fibrils observed here could result from the non-prion like sequence of the FXR2P LC domain.

Our results reveal a context-dependent interplay between RNA binding and LC sequences in regulating the structural state of RBPs in neurons. While FXR2P fibrillogenesis can result from manipulating a short stretch of the LC domain (EGFP insertion within residues 416-439), we found that disrupting the KH2 RNA binding domain abolishes these fibrillar assemblages (Fig. 7J-O). Interestingly, we also observed that ribosomes associate with both isolated Type A and B fibrils (Fig. 4) as well as with bundled Type A fibrils, but Type B bundles are devoid of protein synthesis machinery (Figs. 4 and 6). Notably, ribosomes are only detected in ~50% of FXGs, endogenous FXR2P-containing assemblages (Akins et al., 2017 and see below). Taken together with our observations here, these data suggest that changes in the structural state of FXR2P might influence the ability of FXR2P-containing assemblages such as FXGs to associate with ribosomes and thereby regulate local neuronal translation.

Our work suggests that the FXR2P LC and RNA binding domains could play an important role in the circuit-selective formation of FXGs in the intact brain. FXGs are only present in select axons, such as corticocortical and thalamocortical fibers, olfactory sensory neurons, hippocampal CA3 associational axons and cerebellar parallel fibers (Christie et al., 2009; Akins et al., 2012). Moreover, four FXG subtypes, all containing FXR2P but differing in FMRP/FXR1P and mRNA content, are present in distinct circuits (Christie et al., 2009; Akins et al., 2017; Chyung et al., 2018). These results suggest that individual neuronal types contain sets of molecular and/or cellular factors that promote the structural reorganization of FXR2P via its LC and RNA binding domains to assemble FXGs in a circuit-selective fashion. Elucidation of the factors promoting FXR2P fibrillization and determining if these are regulated in a cell autonomous fashion could provide important insight into how FXGs and their RNA association are regulated in neuronal subsets in the brain.

Finally, these findings have implications for understanding the role of RNA binding proteins in neurological disease. Pathological RNA-protein aggregates caused by RBP dysregulation characterize many neurodegenerative diseases including ALS and FTD (Conlon and Manley 2017; Bowden and Dormann 2016). Moreover, familial forms of these diseases are often caused by LC domain mutations (Purice and Taylor 2018). Such mutations are thought to result in dysregulation of endogenous mechanisms controlling higher order structural states of key RBPs (Kim et al., 2013; Ramaswami et al., 2013; Purice and Taylor 2018). The multiple higher order states of FXR2P defined here could thus shed light on structural perturbations mediating abnormal RBP function in neurodegeneration.

## MATERIALS AND METHODS

### Plasmids

Full-length *fxr2* was PCR generated from an *fxr2* clone (Open Biosystems; MMM1013-9498022) and restriction subcloned into the pBluescriptKSII vector. Construction of the modified pCAG plasmid pCAGES and pCAGES-TdTomato has been described previously (Stackpole et al., 2014). The modified Tn5 transposon encoding *EGFP*-*Kan^R^-STOP* (pBNJ24.6) was a kind gift of Dr. Thomas Hughes (Sheridan et al., 2002). The FXR2P^[217-∆LC]^ plasmid was generated by PCR from the FXR2P^[217]^ clone using the forward primer 5’– GAATTCGATGGGCGGCCTGGCC-3’ and reverse primer 3’-CTCGAGTTAAAAGCCCAGCCCAATCTG-5’. The I314N point mutation was generated in FXR2P^[217]^, FXR2P^[416]^ and FXR2P^[435]^ constructs using GeneArt technologies (ThermoFisher; Grand Island, NY) with the targeted mutation 5’-GTTAAC-3’ which also introduces an HpaI restriction site for selection during cloning.

### In vitro transposition reaction

Transposons were amplified from the pBNJ24.6 plasmid by PCR with a primer complimentary to the Tn5 mosaic end (5’-CTGTCTCTTATACACATCT −3’). The *in vitro* transposition reaction was performed with the amplified transposon and target plasmid (pBluescriptKSII-*fxr2*) using EZ-Tn5 Transposase according to manufacturer’s instructions (Epicentre; Madison, WI). Electrocompetent *E. coli* were transformed with the transposition reaction and plated on LB agar with Kanamycin (30μg/mL) and Ampicillin (100μg/mL). To establish transposition efficiency transformation reactions were plated in parallel on LB agar with Ampicillin alone.

### Generation of Full-Length EGFP-Transposed FXR2 (FXR2P^EGFP^) Constructs

To determine which clones harbored the EGFP transposon insert both in-frame and in the correct orientation within pBluescriptKSII-fxr2, colonies were visually screened for EGFP fluorescence using an Olympus SZX12 microscope. A total of 180 fluorescent colonies were selected and DNA was prepared from each clone (miniprep kits from Qiagen; Valencia, CA). Each fluorescent transposed clone was then screened for insertion of EGFP into the *fxr2* coding region (versus vector backbone) by restriction digestion with XbaI and XhoI. Of these, the exact insertion site of the transposon within *fxr2* was identified by sequencing 5’ out of the transposon using a primer complimentary to the EGFP coding region (3’-TTTACGTCGCCGTCCAGCTCG A-5’). To generate full-length fusion proteins, the Kanamycin selection cassette with *STOP* codon was first removed from all clones with unique in-frame insertion sites by digestion with SrfI (Agilent; Santa Clara, CA) and re-ligation with T4 DNA Ligase (NEB, Ipswich, MA). To verify loss of cassette, all colonies were restriction digested with XmaI and KpnI. Each EGFP-transposed *fxr2* construct was then restriction subcloned into the EcoRV and NheI sites of the pCAGES vector to generate pCAGES-FXR2P^EGFP^ constructs.

### Primary Rat Cortical Neuron Culture

All animal care and collection of tissue were in accordance with Brown University IACUC guidelines for care and use of laboratory animals. Primary rat cortical neuron cultures were prepared as previously described (Stackpole et al., 2014). For microscopy experiments, cells were plated onto 24-well plates with poly-D-lysine (PDL; 50μg/mL; Sigma; St. Louis, MO) and laminin (20μg/mL; Thermo Scientific; Grand Island, NY) coated glass coverslips (1mm; Assistent; Germany) at a density of 80,000 cells/well. For Western blotting, cells were plated onto PDL-coated 6-well plates at a density of 300,000 cells/well. For electron microscopy, cells were plated onto PDL-coated 4-well Permanox Lab-Tek chamberslides at a density of 80,000 cells/well. Cultures were maintained in an incubator at 37˚C and 5% CO_2_ and 95% air.

### Transfection of Primary Cultures

At 3 days *in vitro* (DIV), cultures were co-transfected with pCAGES-FXR2P^EGFP^ constructs along with pCAGES-TdTomato by magnetofection with NeuroMag paramagnetic nanobeads (Oz Biosciences; France). For 24-well plates, plasmid DNA (0.5 μg total, or 0.25 μg each construct) was incubated with 1.75 μL NeuroMag beads for 15 minutes. For 6-well plates (western blotting), 0.75 µg of plasmid DNA was incubated with 2.62 µL of NeuroMag beads. Solution was then added drop-wise to cultures and allowed to incubate for 15 minutes on top a magnetic plate within a 37˚ incubator.

### Immunostaining

At various timepoints post transfection, coverslips were washed once with PBS and fixed for 15 minutes with 4% paraformaldehyde with 4% sucrose in PBS. Coverslips were blocked with PBS with 0.3% Triton X-100 and 1% blocking reagent (Roche; United States) for 30 min and then incubated for 1 hour each in the same solution with primary antibody followed, after washing, by secondary antibodies (Thermo Scientific, Grand Island, NY; 1:1000). Primary antibodies: rRNA 5S and 5.8S subunits were detected using supernatant from hybridoma cells expressing the monoclonal antibody Y10b (gift from Dr. J. Twiss; 1:200). GFP was detected using antibody N86/8 from NeuroMab (Davis, CA; 1:10). Coverslips were mounted in NPG (4% n-propyl-gallate, 60% glycerol, 5 mM phosphate pH 7.4, 75 mM sodium chloride). Confocal images were collected on a Zeiss LSM 510 microscope using z-stacks to capture the entire depth of the neuron using a 63X Plan-Apochromat objective. Images were analyzed using ImageJ and Photoshop CS6 (Adobe, San Jose, CA). To depict structures that existed across a wide dynamic range, nonlinear adjustments were made to brightness and contrast in order to accentuate signal while maintaining background. Quantifications were performed only on images for which linear manipulations were applied uniformly across all images in the dataset regardless of condition.

### In Situ Hybridization

Cultured neurons were washed and fixed as described above and then treated with 0.2M hydrochloric acid for 10 minutes and then PBS with 1% Triton X-100 for 2 min. Cultures were rinsed, equilibriated in 2X SSC with 10% formamide, and incubated overnight at 37˚ in hybridization solution (10% dextran sulfate, 2 mM vanadyl ribonucleosides [NEB; Ipswich, MA], 2X SSC, 10% deionized formamide, 1 mg/mL E. coli tRNA [Roche; United States], 200 µg/mL BSA [Roche; United States]) with oligo(dT)_45_ that had been end-labeled with the DIG oligonucleotide tailing kit following manufacturer’s instructions for short tails (Roche; United States). Coverslips were washed with 2X SSC and 10% formamide for 30 minutes each at 37˚ and rinsed with 2X SSC and PBST. A primary antibody against digoxigenin (1:100, Jackson Immunolabs; West Grove, PA) was applied to cells in blocking solution for 2 h. Cells were then rinsed with PBST, incubated in secondary antibody for 1 h, washed and mounted in NPG medium and analyzed as described above.

### Bioinformatic Predictors of LC Domain Properties

The FXR2P amino acid sequence was assessed for intrinsically disordered regions using the PONDR-FIT website interface (http://www.disprot.org/pondr-fit.php). For prion-like analyses, the FXR2P amino acid sequence was uploaded to the Prion-Like Amino Acid Composition website (PLAAC; http://plaac.wi.mit.edu/) and analyzed with the default settings. The PLAAC website simultaneously includes analyses using FoldIndex and the 4*PAPA algorithm. For predicting disordered binding regions, the FXR2P amino acid sequence was assessed by ANCHOR using the IUPRED website (http://iupred.enzim.hu/). For above analyses, the graphical output from each website was used for display purposes in this paper. The pi-contact propensity of FXR2P was evaluated using a pi-pi phase separation propensity script (DOI: 10.7554/eLife.31486). The per-residue score of the pi-contact propensity was then plotted in MATLAB. The fibril forming propensity of FXR2P was evaluated using the ZipperDB website interface (https://services.mbi.ucla.edu/zipperdb/) and the Rosetta Energy score was then plotted in MATLAB for display purposes.

### Electron Microscopy

DIV6 (FXR2P^[217]^ and ^[416]^) or DIV14 (FXR2P^[435]^) transfected neurons in 4-well chamberslides were washed three times with 1.25% glutaraldehyde in 0.15M sodium cacodylate buffer and fixed overnight at 4˚. A circular diamond scribe objective (Zeiss) was used to score the location of GFP+ transfected neurons (n= 3 per condition) on the bottom of the chamberslide. Epifluorescent images of each identified neuron were collected with a 10X objective using a Zeiss Axiovert 200M microscope to identify the position of transfected neuron in the scored location. Slides were then rinsed and post-fixed with 1% osmium tetroxide, rinsed and dehydrated through a graded ethanol series. After removal of media chambers and gaskets slides were covered with Epox 812 resin, placed over resin filled slide-duplicating molds and polymerized overnight. Regions of interest determined from the epiflourescent images were cut out and mounted. Ultrathin sections (50nm) were collected on wire mesh copper grids incubated with uranyl acetate and lead citrate and examined with a Morgagni 268 transmission electron microscope. Images were collected with an AMT Advantage 542 CCD camera system. The width of isolated fibrils was quantified in ImageJ by measuring the shortest distance across the fibrils perpendicular to the long axis. Depending on the continuous length of the fibrils within individual micrographs, the width was measured and averaged across one to eight sites to give the average diameter of each fibril. Ribosome periodicity was determined in ImageJ by measuring the distance between the centers of adjacent ribosomes along fibrils. Depending on the continuous length of fibrils within individual micrographs, the inter-ribosome distance was measured and averaged across one to six pairs to give the ribosome periodicity for each fibril. Statistics including mean and standard error were calculated in Prism.

### Western Blotting

Cultured cortical neurons (DIV3) were transfected as described above. Three days post transfection (DIV6), neurons were washed twice with cold PBS and incubated with RIPA lysis buffer (25 mM Tris pH 8.0, 150 mM sodium chloride, 0.1% SDS, 1% NP-40, 0.5% sodium deoxycholate, 5 mM EDTA, 2 mM sodium orthovanadate, NEB; Ipswich, MA; 10 mM β-glycerophosphate, 1X protease inhibitor cocktail, Roche; United States) for 15 min at 4°C. Lysates were then collected and centrifuged at 10,000*g* for 15 min at 4°C. Supernatants (10 µg) were boiled in SDS-PAGE sample buffer (60 mM Tris pH 6.8, 2% SDS, 10% glycerol, 735 mM β-mercaptoethanol), separated by SDS-PAGE gels and transferred to nitrocellulose membranes following standard methods. Blots were blocked for 1 h at room temperature in 5% nonfat dried milk and 4% normal goat serum in wash buffer (100 mM Tris pH 7.4, 150 mM sodium chloride, and 0.1% Tween-20) and then incubated overnight with diluted primaries at 4°C (anti-FXR2P clone 55; 1:1000; BD Biosciences; or anti-γ-actin; 1:40,000, Sigma). Blots were washed four times for 5 min each in wash buffer after primary incubation. Horseradish peroxidase (HRP)-conjugated secondary antibodies (KPL, Gaithersburg, MD) were diluted in 5% milk in wash buffer (1:2,000) and incubated at room temperature for 90 min. Blots were washed four times for 5 min each in wash buffer and signals were detected by chemiluminescence (Amersham ECL Western blotting reagents, Arlington Heights, IL).

### Polyribosome Analysis

HEK293T cells were cultured in 150-mm culture dishes and transfected with FXR2P^[217]^ or FXR2P^[217::I314N]^ constructs using the calcium chloride method. After 24 h, cells were harvested by replacing the culture media with fresh media containing cycloheximide (Sigma; St. Louis, MO) at a final concentration of 100 μg/mL for 15 min. Cells were washed twice with ice cold PBS containing 100 μg/mL cycloheximide, trypsinized and pelleted for 5 min at 1000g. Cells were then resuspended in 750μL of low-salt buffer buffer (20 mM Tris-HCl pH 7.5, 10 mM NaCl, 3 mM MgCl_2_) followed by incubation on ice for 5 min. Triton-X 100 was added to the cell suspension to a final concentration of 0.3% (v/v) and cells were lysed on ice using a 1-ml Dounce homogenizer. The solution was centrifuged for 1 min at 10,000*g* at 4 °C, and supernatants were layered on top of linear 15%-50% (w/v) sucrose gradients, ultracentrifuged (Beckman SW41Ti rotor) at 36,000 rpm for 2 h at 4°C. Polysome profiles were monitored by absorbance of light with a wavelength of 254 nm (*A*_254_). For Western analysis, 20μL of each fraction was boiled in SDS-PAGE buffer, separated by SDS-PAGE, transferred to nitrocellulose membrane and then probed by Western analysis for anti-FXR2P (see above).

## Roles of Authors

All authors had full access to all the data in the study and take responsibility for the integrity of the data and the accuracy of the data analysis. Study concept and design: EES, MRA and JRF. EES performed the screening and the characterization studies in neurons. MI performed the sucrose gradients. ACM performed the ZipperDB and pi analyses. NLF assisted with bioinformatic analyses and interpretation. Analysis and interpretation of data: EES, MRA, MI, ACM, NLF and JRF. Drafting of the manuscript: EES and JRF. Critical revision of the manuscript for intellectual content: EES, MRA, and JRF. Obtained funding: MRA and JRF.

## Acknowledgements

We thank C. Ayala, G. Williams, B. McKechnie, V. Medrano, and I. Lopez for technical assistance and J. Richter for technical advice. We thank T. Hughes for the generous gift of the pBNJ24.6 vector. Grant support: HD052083 to JRF and MH090237 to MRA.

## Works Cited

Adams-Cioaba, M.A., Guo, Y., Bian, C., Amaya, M.F., Lam, R., Wasney, G.A., Vedadi, M., Xu, C., and Min, J. (2010). Structural Studies of the Tandem Tudor Domains of Fragile X Mental Retardation Related Proteins FXR1 and FXR2. PLOS ONE 5, e13559.

Akins, M.R., LeBlanc, H.F., Stackpole, E.E., Chyung, E., and Fallon, J.R. (2012). Systematic mapping of Fragile X granules in the developing mouse brain reveals a potential role for presynaptic FMRP in sensorimotor functions. J. Comp. Neurol. 520, 3687–3706.

Akins, M.R., Berk-Rauch, H.E., Kwan, K.Y., Mitchell, M.E., Shepard, K.A., Korsak, L.I.T., Stackpole, E.E., Warner-Schmidt, J.L., Sestan, N., Cameron, H.A., et al. (2017). Axonal ribosomes and mRNAs associate with fragile X granules in adult rodent and human brains. Hum. Mol. Genet. 26, 192–209.

Ascano, M., Mukherjee, N., Bandaru, P., Miller, J.B., Nusbaum, J., Corcoran, D.L., Langlois, C., Munschauer, M., Dewell, S., Hafner, M., et al. (2012). FMR1 targets distinct mRNA sequence elements to regulate protein expression. Nature 492, 382–386.

Batista, A.F.R., Martínez, J.C., and Hengst, U. (2017). Intra-axonal Synthesis of SNAP25 Is Required for the Formation of Presynaptic Terminals. Cell Rep. 20, 3085–3098.

Boeynaems, S., Alberti, S., Fawzi, N.L., Mittag, T., Polymenidou, M., Rousseau, F., Schymkowitz, J., Shorter, J., Wolozin, B., Van Den Bosch, L., et al. (2018). Protein Phase Separation: A New Phase in Cell Biology. Trends Cell Biol. 28, 420–435.

Boke, E., Ruer, M., Wühr, M., Coughlin, M., Lemaitre, R., Gygi, S.P., Alberti, S., Drechsel, D., Hyman, A.A., and Mitchison, T.J. (2016). Amyloid-like Self-assembly of a Cellular Compartment. Cell 166, 637–650.

Bowden, H.A., and Dormann, D. (2016). Altered mRNP granule dynamics in FTLD pathogenesis. J. Neurochem. 138 Suppl 1, 112–133.

Buchan, J.R. (2014). mRNP granules. RNA Biol. 11, 1019–1030.

Burke, K.A., Janke, A.M., Rhine, C.L., and Fawzi, N.L. (2015). Residue-by-Residue View of In Vitro FUS Granules that Bind the C-Terminal Domain of RNA Polymerase II. Mol. Cell 60, 231–241.

Calabretta, S., and Richard, S. (2015). Emerging Roles of Disordered Sequences in RNA-Binding Proteins. Trends Biochem. Sci. 40, 662–672.

Chong, P.A., and Forman-Kay, J.D. (2016). Liquid–liquid phase separation in cellular signaling systems. Curr. Opin. Struct. Biol. 41, 180–186.

Christie, S.B., Akins, M.R., Schwob, J.E., and Fallon, J.R. (2009). The FXG: A Presynaptic Fragile X Granule Expressed in a Subset of Developing Brain Circuits. J. Neurosci. 29, 1514–1524.

Chyung, E., LeBlanc, H.F., Fallon, J.R., and Akins, M.R. (2018). Fragile X granules are a family of axonal ribonucleoprotein particles with circuit-dependent protein composition and mRNA cargos. J. Comp. Neurol. 526, 96–108.

Conlon, E.G., and Manley, J.L. (2017). RNA-binding proteins in neurodegeneration: mechanisms in aggregate. Genes Dev. 31, 1509–1528.

Cumberworth, A., Lamour, G., Babu, M.M., and Gsponer, J. (2013). Promiscuity as a functional trait: intrinsically disordered regions as central players of interactomes. Biochem. J. 454, 361–369.

Darnell, R.B. (2013). RNA Protein Interaction in Neurons. Annu. Rev. Neurosci. 36, 243–270.

Darnell, J.C., Fraser, C.E., Mostovetsky, O., Stefani, G., Jones, T.A., Eddy, S.R., and Darnell, R.B. (2005a). Kissing complex RNAs mediate interaction between the Fragile-X mental retardation protein KH2 domain and brain polyribosomes. Genes Dev. 19, 903–918.

Darnell, J.C., Mostovetsky, O., and Darnell, R.B. (2005b). FMRP RNA targets: identification and validation. Genes Brain Behav. 4, 341–349.

Darnell, J.C., Fraser, C.E., Mostovetsky, O., and Darnell, R.B. (2009). Discrimination of common and unique RNA-binding activities among Fragile X mental retardation protein paralogs. Hum. Mol. Genet. 18, 3164–3177.

Dosztányi, Z., Mészáros, B., and Simon, I. (2009). ANCHOR: web server for predicting protein binding regions in disordered proteins. Bioinformatics 25, 2745–2746.

Dunker, A.K., Brown, C.J., Lawson, J.D., Iakoucheva, L.M., and Obradović, Z. (2002). Intrinsic disorder and protein function. Biochemistry (Mosc.) 41, 6573–6582.

Elbaum-Garfinkle, S., Kim, Y., Szczepaniak, K., Chen, C.C.-H., Eckmann, C.R., Myong, S., and Brangwynne, C.P. (2015). The disordered P granule protein LAF-1 drives phase separation into droplets with tunable viscosity and dynamics. Proc. Natl. Acad. Sci. 112, 7189–7194.

Feng, Y., Absher, D., Eberhart, D.E., Brown, V., Malter, H.E., and Warren, S.T. (1997). FMRP Associates with Polyribosomes as an mRNP, and the I304N Mutation of Severe Fragile X Syndrome Abolishes This Association. Mol. Cell 1, 109–118.

Giraldez, T., Hughes, T.E., and Sigworth, F.J. (2005). Generation of Functional Fluorescent BK Channels by Random Insertion of GFP Variants. J. Gen. Physiol. 126, 429–438.

Goldschmidt, L., Teng, P.K., Riek, R., and Eisenberg, D. (2010). Identifying the amylome, proteins capable of forming amyloid-like fibrils. Proc. Natl. Acad. Sci. 107, 3487–3492.

Guo, L., and Shorter, J. (2015). It’s Raining Liquids: RNA Tunes Viscoelasticity and Dynamics of Membraneless Organelles. Mol. Cell 60, 189–192.

Han, T.W., Kato, M., Xie, S., Wu, L.C., Mirzaei, H., Pei, J., Chen, M., Xie, Y., Allen, J., Xiao, G., et al. (2012). Cell-free Formation of RNA Granules: Bound RNAs Identify Features and Components of Cellular Assemblies. Cell 149, 768–779.

Harrison, A.F., and Shorter, J. (2017). RNA-binding proteins with prion-like domains in health and disease. Biochem. J. 474, 1417–1438.

Holt, C.E., and Schuman, E.M. (2013). The Central Dogma Decentralized: New Perspectives on RNA Function and Local Translation in Neurons. Neuron 80, 648–657.

Kapeli, K., and Yeo, G. (2012). Genome-Wide Approaches to Dissect the Roles of RNA Binding Proteins in Translational Control: Implications for Neurological Diseases. Front. Neurosci. 6.

Kapeli, K., Martinez, F.J., and Yeo, G.W. (2017). Genetic mutations in RNA-binding proteins and their roles in ALS. Hum. Genet. 136, 1193–1214.

Kato, M., Han, T.W., Xie, S., Shi, K., Du, X., Wu, L.C., Mirzaei, H., Goldsmith, E.J., Longgood, J., Pei, J., et al. (2012). Cell-free Formation of RNA Granules: Low Complexity Sequence Domains Form Dynamic Fibers within Hydrogels. Cell 149, 753–767.

Kiebler, M.A., and Bassell, G.J. (2006). Neuronal RNA Granules: Movers and Makers. Neuron 51, 685–690.

Kim, H.J., Kim, N.C., Wang, Y.-D., Scarborough, E.A., Moore, J., Diaz, Z., MacLea, K.S., Freibaum, B., Li, S., Molliex, A., et al. (2013). Mutations in prion-like domains in hnRNPA2B1 and hnRNPA1 cause multisystem proteinopathy and ALS. Nature 495, 467–473.

King, O.D., Gitler, A.D., and Shorter, J. (2012). The tip of the iceberg: RNA-binding proteins with prion-like domains in neurodegenerative disease. Brain Res. 1462, 61–80.

Korsak, L.I.T., Mitchell, M.E., Shepard, K.A., and Akins, M.R. (2016). Regulation of neuronal gene expression by local axonal translation. Curr. Genet. Med. Rep. 4, 16–25.

Laggerbauer, B., Ostareck, D., Keidel, E.M., Ostareck-Lederer, A., and Fischer, U. (2001). Evidence that fragile X mental retardation protein is a negative regulator of translation. Hum. Mol. Genet. 10, 329–338.

Lancaster, A.K., Nutter-Upham, A., Lindquist, S., and King, O.D. (2014). PLAAC: a web and command-line application to identify proteins with prion-like amino acid composition. Bioinformatics 30, 2501–2502.

Levenga, J., Buijsen, R.A.M., Rifé, M., Moine, H., Nelson, D.L., Oostra, B.A., Willemsen, R., and de Vrij, F.M.S. (2009). Ultrastructural analysis of the functional domains in FMRP using primary hippocampal mouse neurons. Neurobiol. Dis. 35, 241–250.

Mackenzie, I.R., Nicholson, A.M., Sarkar, M., Messing, J., Purice, M.D., Pottier, C., Annu, K., Baker, M., Perkerson, R.B., Kurti, A., et al. (2017). TIA1 Mutations in Amyotrophic Lateral Sclerosis and Frontotemporal Dementia Promote Phase Separation and Alter Stress Granule Dynamics. Neuron 95, 808–816.e9.

Maziuk, B., Ballance, H.I., and Wolozin, B. (2017). Dysregulation of RNA Binding Protein Aggregation in Neurodegenerative Disorders. Front. Mol. Neurosci. 10.

Molliex, A., Temirov, J., Lee, J., Coughlin, M., Kanagaraj, A.P., Kim, H.J., Mittag, T., and Taylor, J.P. (2015). Phase Separation by Low Complexity Domains Promotes Stress Granule Assembly and Drives Pathological Fibrillization. Cell 163, 123–133.

Mészáros, B., Simon, I., and Dosztányi, Z. (2009). Prediction of Protein Binding Regions in Disordered Proteins. PLOS Comput. Biol. 5, e1000376.

Mitrea, D.M., and Kriwacki, R.W. (2016). Phase separation in biology; functional organization of a higher order. Cell Commun. Signal. 14, 1.

Murakami, T., Qamar, S., Lin, J.Q., Schierle, G.S.K., Rees, E., Miyashita, A., Costa, A.R., Dodd, R.B., Chan, F.T.S., Michel, C.H., et al. (2015). ALS/FTD Mutation-Induced Phase Transition of FUS Liquid Droplets and Reversible Hydrogels into Irreversible Hydrogels Impairs RNP Granule Function. Neuron 88, 678–690.

Nielsen, F.C., Hansen, H.T., and Christiansen, J. (2016). RNA assemblages orchestrate complex cellular processes. Bioessays 38, 674–681.

Prilusky, J., Felder, C.E., Zeev-Ben-Mordehai, T., Rydberg, E.H., Man, O., Beckmann, J.S., Silman, I., and Sussman, J.L. (2005). FoldIndex©: a simple tool to predict whether a given protein sequence is intrinsically unfolded. Bioinformatics 21, 3435–3438.

Purice, M.D., and Taylor, J.P. (2018). Linking hnRNP Function to ALS and FTD Pathology. Front. Neurosci. 12.

Ramaswami, M., Taylor, J.P., and Parker, R. (2013). Altered Ribostasis: RNA-Protein Granules in Degenerative Disorders. Cell 154, 727–736.

Schluth-Bolard, C., Sanlaville, D., Labalme, A., Till, M., Morin, I., Touraine, R., and Edery, P. (2010). 17p13.1 microdeletion involving the TP53 gene in a boy presenting with mental retardation but no tumor. Am. J. Med. Genet. A. 152A, 1278–1282.

Schwartz, J.C., Wang, X., Podell, E.R., and Cech, T.R. (2013). RNA Seeds Higher-Order Assembly of FUS Protein. Cell Rep. 5, 918–925.

Sheridan, D.L., Berlot, C.H., Robert, A., Inglis, F.M., Jakobsdottir, K.B., Howe, J.R., and Hughes, T.E. (2002). A new way to rapidly create functional, fluorescent fusion proteins: random insertion of GFP with an in vitro transposition reaction. BMC Neurosci. 3, 7.

Shigeoka, T., Jung, H., Jung, J., Turner-Bridger, B., Ohk, J., Lin, J.Q., Amieux, P.S., and Holt, C.E. (2016). Dynamic Axonal Translation in Developing and Mature Visual Circuits. Cell 166, 181–192.

Stackpole, E.E., Akins, M.R., and Fallon, J.R. (2014). N-myristoylation regulates the axonal distribution of the Fragile X-related protein FXR2P. Mol. Cell. Neurosci. 62, 42–50.

Stepniak, B., Kästner, A., Poggi, G., Mitjans, M., Begemann, M., Hartmann, A., der Auwera, S.V., Sananbenesi, F., Krueger-Burg, D., Matuszko, G., et al. (2015). Accumulated common variants in the broader fragile X gene family modulate autistic phenotypes. EMBO Mol. Med. 7, 1565–1579.

Sutton, M.A., and Schuman, E.M. (2006). Dendritic Protein Synthesis, Synaptic Plasticity, and Memory. Cell 127, 49–58.

Tamanini, F., Bontekoe, C., Bakker, C.E., van Unen, L., Anar, B., Willemsen, R., Yoshida, M., Galjaard, H., Oostra, B.A., and Hoogeveen, A.T. (1999). Different targets for the fragile X-related proteins revealed by their distinct nuclear localizations. Hum. Mol. Genet. 8, 863–869.

Tamanini, F., Kirkpatrick, L.L., Schonkeren, J., van Unen, L., Bontekoe, C., Bakker, C., Nelson, D.L., Galjaard, H., Oostra, B.A., and Hoogeveen, A.T. (2000). The fragile X-related proteins FXR1P and FXR2P contain a functional nucleolar-targeting signal equivalent to the HIV-1 regulatory proteins. Hum. Mol. Genet. 9, 1487–1493.

Taylor, A.M., Berchtold, N.C., Perreau, V.M., Tu, C.H., Jeon, N.L., and Cotman, C.W. (2009). Axonal mRNA in Uninjured and Regenerating Cortical Mammalian Axons. J. Neurosci. 29, 4697–4707.

Tompa, P. (2012). Intrinsically disordered proteins: a 10-year recap. Trends Biochem. Sci. 37, 509–516.

Toombs, J.A., Petri, M., Paul, K.R., Kan, G.Y., Ben-Hur, A., and Ross, E.D. (2012). De novo design of synthetic prion domains. Proc. Natl. Acad. Sci. 109, 6519–6524.

Toretsky, J.A., and Wright, P.E. (2014). Assemblages: Functional units formed by cellular phase separation. J Cell Biol 206, 579–588.

Vernon, R.M., Chong, P.A., Tsang, B., Kim, T.H., Bah, A., Farber, P., Lin, H., and Forman-Kay, J.D. (2018). Pi-Pi contacts are an overlooked protein feature relevant to phase separation.

Wang, E.T., Taliaferro, J.M., Lee, J.-A., Sudhakaran, I.P., Rossoll, W., Gross, C., Moss, K.R., and Bassell, G.J. (2016). Dysregulation of mRNA Localization and Translation in Genetic Disease. J. Neurosci. 36, 11418–11426.

Weber, S.C., and Brangwynne, C.P. (2012). Getting RNA and Protein in Phase. Cell 149, 1188–1191.

Xiang, S., Kato, M., Wu, L.C., Lin, Y., Ding, M., Zhang, Y., Yu, Y., and McKnight, S.L. (2015). The LC Domain of hnRNPA2 Adopts Similar Conformations in Hydrogel Polymers, Liquid-like Droplets, and Nuclei. Cell 163, 829–839.

Xue, B., Dunbrack, R.L., Williams, R.W., Dunker, A.K., and Uversky, V.N. (2010). PONDR-FIT: A Meta-Predictor of Intrinsically Disordered Amino Acids. Biochim. Biophys. Acta 1804, 996–1010.

Liu-Yesucevitz, L., Bassell, G.J., Gitler, A.D., Hart, A.C., Klann, E., Richter, J.D., Warren, S.T., and Wolozin, B. (2011). Local RNA Translation at the Synapse and in Disease. J. Neurosci. 31, 16086–16093.

Zang, J.B., Nosyreva, E.D., Spencer, C.M., Volk, L.J., Musunuru, K., Zhong, R., Stone, E.F., Yuva-Paylor, L.A., Huber, K.M., Paylor, R., et al. (2009). A Mouse Model of the Human Fragile X Syndrome I304N Mutation. PLoS Genet 5, e1000758.

Zhang, Y., O’Connor, J.P., Siomi, M.C., Srinivasan, S., Dutra, A., Nussbaum, R.L., and Dreyfuss, G. (1995). The fragile X mental retardation syndrome protein interacts with novel homologs FXR1 and FXR2. EMBO J. 14, 5358–5366.

Zheng, J.-Q., Kelly, T.K., Chang, B., Ryazantsev, S., Rajasekaran, A.K., Martin, K.C., and Twiss, J.L. (2001). A Functional Role for Intra-Axonal Protein Synthesis during Axonal Regeneration from Adult Sensory Neurons. J. Neurosci. 21, 9291–9303.

